# The role of trade-offs and feedbacks in shaping integrated plasticity and behavioral correlations

**DOI:** 10.1101/2021.07.26.453877

**Authors:** Ned A. Dochtermann

## Abstract

How behaviors vary among individuals and covary with other behaviors has been a major topic of interest over the last two decades. Unfortunately, proposed theoretical and conceptual frameworks explaining the seemingly ubiquitous observation of behavioral (co)variation have rarely successfully generalized. Two observations perhaps explain this failure: First, phenotypic correlations between behaviors are more strongly influenced by correlated and reversible plastic changes in behavior than by “behavioral syndromes”. Second, while trait correlations are frequently assumed to arise via trade-offs, the observed pattern of correlations is not consistent with simple pair-wise trade-offs. A possible resolution to the apparent inconsistency between observed correlations and a role for trade-offs is provided by state-behavior feedbacks. This is critical because the inconsistency between data and theory represents a major failure in our understanding of behavioral evolution. These two primary observations emphasize the importance of an increased research focus on correlated reversible plasticity in behavior—frequently estimated and then disregarded as within-individual covariances.

**LAY SUMMARY:** Correlations between behaviors are common but observed patterns of these correlations are, at least superficially, inconsistent with expectations of trade-offs. This mismatch is potentially resolved via feedbacks between behaviors and energy availability, suggesting important new research directions.

**DATA AND MODEL AVAILABILITY:** Model code, as well as the data associated with Figures 2 & 3, are available at github.com/DochtermannLab/FeedbacksModel. Both code and data will be made available at Dryad if accepted.

## INTRODUCTION

Despite almost two decades of intensive research effort, overarching explanations for why behavioral responses are frequently correlated remain elusive. Here, I argue that this stems from two key empirical findings that have not been fully appreciated. First, behavioral correlations are more strongly influenced by within-individual correlations than they are by behavioral syndromes (Table 1A, Box1). Second, bivariate trade-offs are an insufficient explanation for—and inconsistent with—observed behavioral correlations. Instead, observed correlations are consistent with a combination of feedbacks (positive and negative) between behaviors and states.

**Table 1.** A. Parameters characterizing behavioral associations. B. Some possible contributors to within-individual behavioral correlations. If either active or passive plasticity are elicited in response to known and measured environmental parameters, they can be explicitly modeled. Otherwise, they will contribute to within-individual variation and covariation. Similar processes contribute to the irreversible plasticity (aka developmental plasticity), which contributes to among-individual correlations (Dingemanse and Dochtermann 2014). The contribution of each component of within-individual correlations can be determined by breaking *r*_W_ down into constituent parts, similar to how a phenotypic correlation was broken down in Box 1.

| A. | Also known as | Definition |  |
| --- | --- | --- | --- |
| Phenotypic correlations |  | The standardized covariance between two behaviors. Describes the strength and direction of association. | (Sih et al. 2004a, Sih et al. 2004b, Dingemanse et al. 2010) |
| Among-individual correlations | Behavioral syndromes | Correlations in an individual's behavioral responses | (Dingemanse et al. 2012, Dingemanse and Dochtermann 2013) |
| Within-individual correlations | Residual correlations | Correlations in *changes* in an individual's responses |  |
| B. | Definition |  |  |
| Active reversible plasticity | changes in an individual's behavior expressed in response to environmental cues indicative of selective pressures |  | (Piersma and Drent 2003, Piersma and Van Gils 2011) |
| Passive reversible plasticity | changes in response to environmental conditions rather than specific cues of selective pressures; includes passive responses to abiotic conditions, such as hypoxia |  | (Whitman and Agrawal 2009) |
| Reversible organismal error | changes to an organism's phenotype in response to an incorrectly processed cue |  | (Westneat et al. 2015) |
| Measurement error | error in quantification of behaviors; can be correlated due to bias and |  | (Westneat et al. 2015) |

for a discussion of additional contributors to within-individual (co)variation see: (Piersma and Drent 2003, Whitman and Agrawal 2009, Piersma and Van Gils 2011, Westneat et al. 2015, Berdal and Dochtermann 2019)

### 1. Understanding how among and within individual variation contribute to behavioral correlations

Much of the research examining behavioral correlations over the last two decades has focused on among-individual correlations, i.e., “behavioral syndromes” (Figure 1A & 1B, Box 1; Sih et al. 2004a, Sih et al. 2004b, Dingemanse et al. 2010, Dingemanse et al. 2012). This interest has been reasonable because behavioral syndromes connect to both underlying genetic correlations and developmental processes. As such, behavior correlations can directly influence evolutionary outcomes (Dochtermann and Dingemanse 2013, Royauté et al. 2020). This clearly embeds behavioral syndrome research into the broader realms of evolutionary biology. Unfortunately, this focus on syndromes ignores the fact that behavioral correlations are, instead, more strongly influenced by within-individual correlations (Box 1, Figure 1, Table 1A). This has led to a disconnect between research effort and biology.

**Figure 1.**
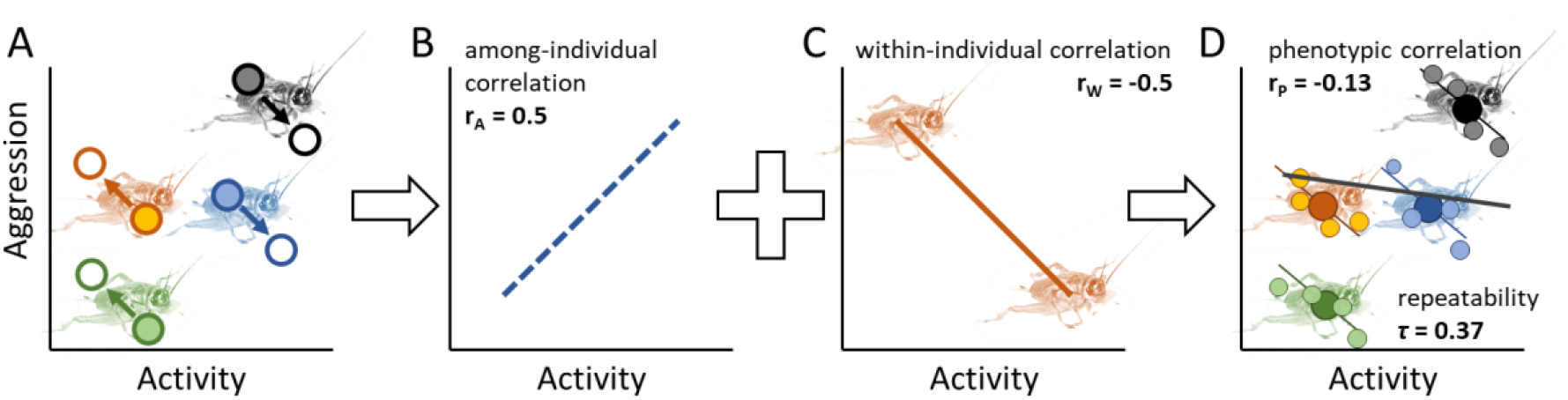
Example of how behavioral syndromes, i.e. among-individual correlations (*r*_A_) and correlated plasticity, i.e. within-individual correlations (*r*_W_), additively contribute to phenotypic correlations. A) Activity and aggressive behavior of four crickets are measured on two occasions (1 & 2). Crickets largely maintain their relative differences in each behavior (i.e. “personality”) *and* their behavior changes in a correlated manner (correlated plasticity). B) The relationship *among* crickets is due to a positive among-individual correlation (dashed blue line, *r*_A_). C) Behavioral plasticity is negatively correlated (within-individual correlation: solid orange line, *r*_W_): if an *individual* increases its activity, it decreases its aggression. D) The among- and within-individual correlations additively determine average individual behaviors (dark circles) and how crickets vary from their averages (colored lines and smaller, lighter, circles). The influence of either *r*_A_ or *r*_W_ on the phenotypic correlation (solid black line in D) is determined by the repeatabilities of the behaviors. If repeatability is high (> 0.5), this phenotypic correlation is primarily determined by the behavioral syndrome, *rA* (Box 1). Here, with a repeatability of 0.37 (Bell et al. 2009), the phenotypic correlation is more strongly influenced by the within-individual correlation than the among-individual correlation but is ultimately quite modest.

Within-individual correlations represent the occurrence of correlated *changes* in behaviors at the level of an individual. For example, a negative within-individual correlation between aggression and exploratory propensity would mean that when an individual increases its aggression, it also decreases its activity (Figure 1). This correlated change can occur even if, on average, more aggressive individuals are also more active (Figure 1).

That behavioral correlations at the phenotypic level are more strongly influenced by within-individual correlations than by behavioral syndromes is a necessary conclusion given observed behavioral repeatabilities. Due to the mathematical relationship of a phenotypic correlation to its constituent parts (Box 1), whenever repeatability is less than 0.5, the phenotypic correlation is more strongly influenced by within-individual correlations than among-individual correlations (Dingemanse et al. 2012, Dingemanse and Dochtermann 2013). Bell et al.’s (2009) meta-analysis of repeatabilities—a keystone contribution to our understanding of behavioral variation—found that the average repeatability of behaviors was about 0.4. Consequently, behavioral correlations are, on average, most strongly influenced by within-individual correlations (Box 1).

Within-individual correlations include—but are not limited to (see Table 1B)— correlated plastic responses to temporary environmental variation; specifically, responses to unmeasured or unmodeled environmental variation. This correlated reversible plasticity is akin to “phenotypic flexibility” (Piersma and Van Gils 2011) but correlated across multiple behaviors. Despite more strongly influencing behavioral correlations than do behavioral syndromes (Box 1), within-individual correlations have received much less direct theoretical or empirical attention from behavioral researchers.

#### Box 1.

The relationship of behavioral syndromes and reversible plasticity to behavioral correlations: the primacy of correlated reversible plasticity

Behavioral correlations at the population and phenotypic level (*r*_p_) emerge from the joint contributions of among-individual correlations, i.e. behavioral syndromes (*r*_A_), and within-individual correlations (*r*_w_, Figure 1; (Dingemanse et al. 2012)). The relative influence of each of these on observed correlations is mediated by the repeatability of the behaviors of interest (*τ*_1_ and *τ*_2_):

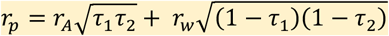

*r*_A_ is the statistical definition of a behavioral syndrome (Dingemanse et al. 2012) and, as repeatability decreases, the contribution of behavioral syndromes to phenotypic correlations necessarily decreases. At the extreme, if *τ* for either behavior is 0, then *r*_p_ is entirely determined by the *r*_w_. Moreover, *r*_w_ contributes more strongly to *r*_p_ than does *r*_A_ when geometric mean repeatability is less than 0.5 (Dingemanse and Dochtermann 2013). Based on meta-analysis (Bell et al. 2009) we know that average *τ*s are around 0.4 and, consequently, *r*_w_, on average influences *r*_p_ more strongly than does *r*_A_. Put another way, within-individual correlations have an average 1.5 times greater influence on phenotypic correlations than do behavioral syndromes.

Importantly, *r*_*w*_ includes correlated reversible plasticity (Figure 1), though this plasticity will not necessarily be adaptive and errors both in the response to cues by organisms or in measurement also contribute to *r*_*w*_ (Table 1).

### 2. Interpreting patterns of correlations and indications, or lack thereof, of a role for trade-offs

A possible factor shaping correlated plastic responses are trade-offs in investment of energy (or other resources) into behaviors. For example, energy used during agonistic interactions is not available for exploring new areas. Accordingly, we would expect that as individuals increase investment in one behavior, they decrease investment in another (Figure 1A), generating negative within-individual correlations (Figure 1C). If individuals differ in their ability to acquire resources, this results in the familiar “big house, big car” scenario from life-history theory (Van Noordwijk and de Jong 1986, Reznick et al. 2000): high variation in acquisition (relative to variation in allocation) results in a positive among-individual correlation despite underlying trade-offs. However, because behavioral researchers frequently change the sign of behavioral measures to aid in the ecological interpretation of behavioral assays, it is not the case that among-individual correlations are strictly expected to be positive and within-individual correlations be negative. Instead, if trade-offs underpin behavioral correlations, we would expect among-individual and within-individual correlations to be of *opposite signs* (Figure 1D; Downs and Dochtermann 2014).

Meta-analyses have examined behavioral correlations at the genetic, among-individual, and within-individual levels (Dochtermann 2011, Brommer and Class 2017). These analyses have demonstrated that correlations across levels are generally concordant as to their signs and magnitudes (Figure 2). As suggested above, this concordance of signs suggests either a *potential* lack of trade-offs in the expression of behaviors or that variation in acquisition is low relative to variation in allocation. Of course, it is also possible that many of the analyzed behaviors should not be expected to trade-off due to naming fallacies obscuring that the same behavior is actually being measured in different ways (Carter et al. 2013) or because limiting resources differ between behaviors. Moreover, because phenotypes consist of many, many behaviors and other traits, any trade-offs would not typically be expected to exist solely between two traits. Nonetheless, the results in Figure 2 suggest that bivariate trade-offs alone are *insufficient* to explain patterns of among- and within-individual correlations alone.

**Figure 2.**
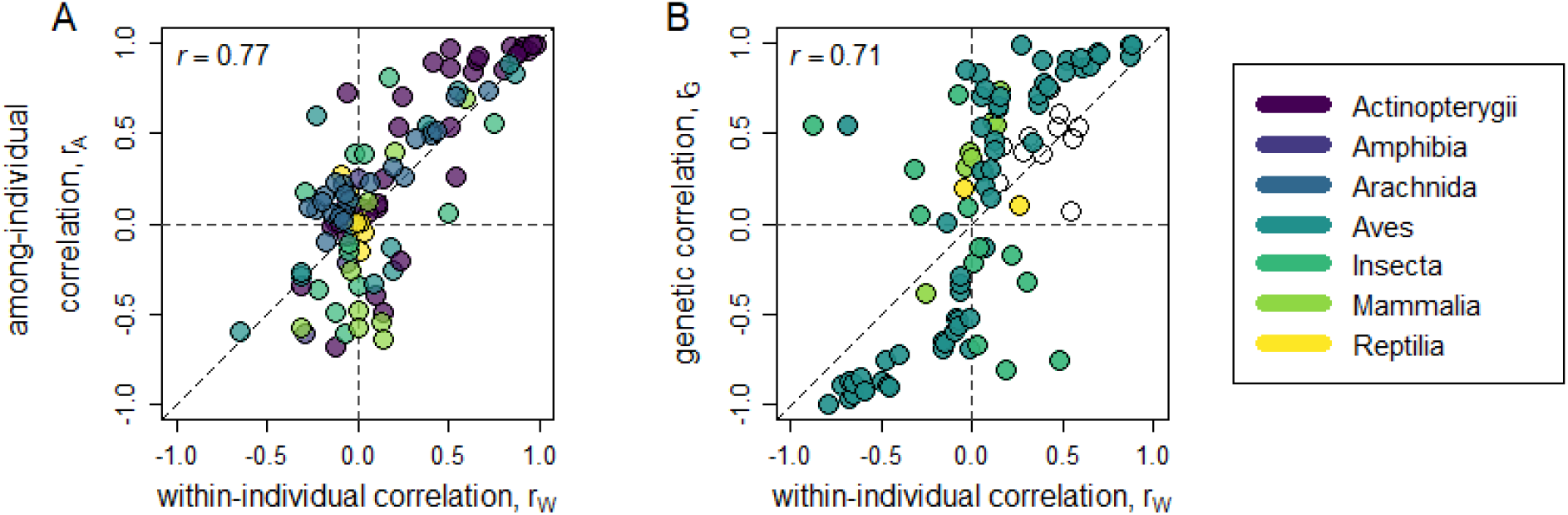
Relationship of within-individual behavioral correlations (*r*_W_) with (A) among-individual correlations (*r*_A_) and (B) genetic correlations (*r*G). The sign and magnitude of *r*_A_ and *r*G are highly concordant with *r*_W_ across behaviors and taxa. Diagonal lines indicate 1:1 relationship. Horizontal and vertical dashed lines divide plots into sign mismatches (top left, bottom right) and sign matches (top right, bottom left). Within-individual correlations were the same sign as among-individual and genetic correlations in 62% and 79% of cases respectively. Data in A are from Brommer and Class (2017). Data in B are from Dochtermann (2011).

### 3. Feedbacks as a potential component influencing behavioral correlations

An alternative mechanism for understanding behavioral correlations builds on a simple model of trade-offs by adding feedbacks between underlying state and behaviors. Sih et al. (2015) proposed that among-individual differences in behavior—i.e., “personality”—can be generated by feedbacks between behavior and underlying state. Specifically, Sih et al. (2015) proposed that positive feedbacks can explain the emergence of personality variation. This was supported by a model where the intensity of initial behavioral expression resulted in feedbacks that then shaped an individual’s future behavioral expression.

To determine whether feedbacks similarly produce both behavioral syndromes and within-individual correlations, I extended the Sih et al. (2015) model to two traits following Houle’s (1991) classic y-model (Figure 3A). The y-model is a simple representation of trade-offs where “state” or some other proxy for energy (S_1_) is allocated to expression of one of two traits (B_1_ or B_2_, Figure 3A). Because energy allocated to one trait at a particular time (TO × S_1,t_) is not available for allocation to the other trait ((1-TO) × S_2,t_), a trade-off exists between them.

**Figure 3.**
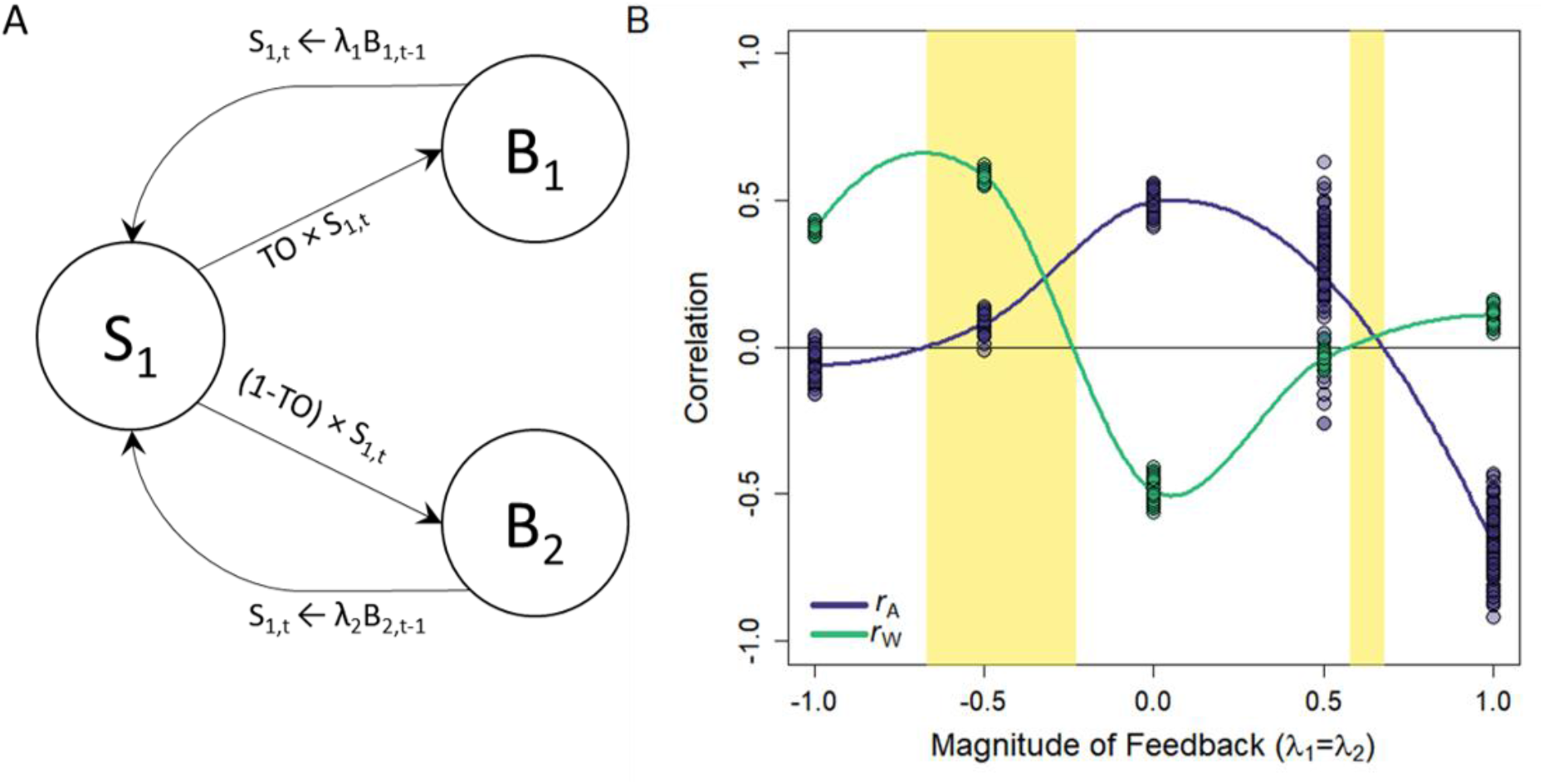
(A) Simple model structure combining Sih et al.’s (2015) feedback model with Houle’s (1991) y-model to include feedbacks between behavior and state. TO (the trade-off parameter) is the proportion of state energy (S1 at time t) allocated to behavior B1, making 1-TO the energy allocated to B2. λ is the feedback strength and the conversion rate of a behavior (B1 or B2) into energy that can be used at a later time. (B) Magnitude of among-(purple) and within-individual (green) correlations under feedbacks (λ) of different strength. Shaded regions indicate those feedback strengths that produce correlations of the same sign. Repeatability is not set *a priori* and is instead an emergent property of the model. Individuals within a population expressed behaviors according to A over ten time steps, with 250 individuals per population. Fifty populations were simulated at each of five levels of feebacks (λ in B). *r*_A_ and *r*_W_ were estimated using the MCMCglmm package in R (Hadfield 2010).

Following this framework, I modeled the expression of two behaviors (B_1_ and B_2_) as emerging from the investment of energy (based on “state”) into each, with the amount of energy invested trading off between behaviors. In this model the two behaviors subsequently affect the gain of state/resources for use in the future expression of behavior via feedbacks (λ, the proportion of energy added back to S_1_, Figure 3A). Biologically, such feedbacks might arise if, for example, exploratory propensity is positively correlated with energy gain. In such a case, individuals with an initially high state can explore more, gaining more energy that can be invested in even greater exploration and energy gain in the future—a positive feedback.

I modeled varied magnitudes of feedbacks, both positive and negative, and analyzed modeled populations to determine resulting the among- and within-individual correlations (Figure 3B). At the start of the model, state differed among individuals due to stochastic variation. This seeded the population with variation in initial acquisition. In natural populations, individuals may similarly initially differ due to developmental stochasticity, parental investment, genetic differences, and other environmental factors.

In the absence of feedbacks (λ_1_ = λ_2_ = 0, Figure 3), this model reduces to the general structure of models developed by Houle (1991) and Van Noordwijk and de Jong (1986). Consequently, without feedbacks, the trade-off in allocation along with underlying variation in initial acquisition results in positive among-individual (*r*_A_) and negative within-individual correlations (*r*_w_, Figure 3B), i.e., negatively correlated plasticity plus different signs for the two correlations. In this case, within-individual variation is generated by stochasticity in how much is invested in one behavior and thus how much is available for the other. For example, individuals might differentially allocate energy to growth versus activity at different periods of their life.

In contrast, with both negative and positive feedbacks (negative and positive values of λ respectively), the within-individual correlation becomes increasingly positive (Figure 3B). Simultaneously, negative and positive feedbacks lead to a reduction in among-individual correlations, with these correlations becoming negative and trade-offs becoming particularly apparent at the among-individual level with strong positive feedback. Importantly, this model identifies both positive and negative feedbacks in two disjunct ranges that result in among-individual correlations and integrated plasticity of the same sign (shaded regions of Figure 3B). This suggests that feedbacks might partially explain the patterns of correlations observed in Figure 2.

The potential for state-behavior feedbacks combined with trade-offs to explain observed patterns of correlations adds to several prior models that suggested state might generate consistent individual differences and phenotypic behavioral correlations (Dall 2004, Wolf et al. 2007, Luttbeg and Sih 2010, Dall et al. 2012). However, these prior models did not separately examine among- and within-individual correlations nor the relationship between them. State-behavior feedbacks are also implied in existing conceptual frameworks, such as the pace-of-life syndrome hypothesis (Réale et al. 2010), though support for such overarching explanations remains elusive (e.g. Niemelä and Dingemanse 2018, Royauté et al. 2018).

### 4. Conclusions

Given that within-individual correlations more strongly influence observed correlations between behaviors than do behavioral syndromes, the mismatch in effort to understand these correlations over the last fifteen years has hindered our general understanding of behavioral correlations. While considerably more investigation is required, the simple model presented here suggests that feedbacks might play a role in shaping both behavioral syndromes and integrated plasticity. This potential for feedbacks to shape correlations can be empirically addressed via manipulation of food availability (e.g. MacGregor et al. 2021), manipulating population densities, and longitudinal measurements of both state and behavior (Sih et al. 2015). Importantly, within-individual correlations can only be estimated with specific study designs (Dingemanse and Dochtermann 2013). Regardless, whether focused on feedbacks or other potential causes, our understanding of behavioral correlations can only advance if more—and careful— attention is paid to within-individual correlations.

## ACKNOWLEDGMENTS

The author thanks B. Hedinger, R. Royauté, J. Saltz, and D. Westneat for helpful discussions. The author also thanks two anonymous for critical and helpful feedback. This feedback improved the clarity and accuracy of the manuscript. This work was funded by NSF-IOS 1557951

